# Could network structures generated with simple rules imposed on a cubic lattice reproduce the structural descriptors of globular proteins?

**DOI:** 10.1101/2020.10.01.321992

**Authors:** Osman Burak Okan, Deniz Turgut, Canan Atilgan, Ali Rana Atilgan, Rahmi Ozisik

## Abstract

A direct way to spot structural features that are universally shared among proteins is to find proper analogues from simpler condensed matter systems. In most cases, sphere-packing arguments provide a straightforward route for structural comparison, as they successfully characterize a wide array of materials such as close packed crystals, dense liquids, and structural glasses. In the current study, the feasibility of creating ensembles of artificial structures that can automatically reproduce a large number of geometrical and topological descriptors of globular proteins is investigated. Towards this aim, a simple cubic (SC) arrangement is shown to provide the best background lattice after a careful analysis of the residue packing trends from 210 proteins. It is shown that a minimalistic set of ground rules imposed on this lattice is sufficient to generate structures that can mimic real proteins. In the proposed method, 210 such structures are generated by randomly removing residues (beads) from clusters that have a SC lattice arrangement until a predetermined residue concentration is achieved. All generated structures are checked for residue connectivity such that a path exists between any two residues. Two additional sets are prepared from the initial structures via random relaxation and a reverse Monte Carlo simulated annealing (RMC-SA) algorithm, which targets the average radial distribution function (RDF) of 210 globular proteins. The initial and relaxed structures are compared to real proteins via RDF, bond orientational order parameters, and several descriptors of network topology. Based on these features, results indicate that the structures generated with 40% occupancy via the proposed method closely resemble real residue networks. The broad correspondence established this way indicates a non-superficial link between the residue networks and the defect laden cubic crystalline order. The presented approach of identifying a minimalistic set of operations performed on a target lattice such that each resulting cluster possess structural characteristics largely indistinguishable from that of a coarse-grained globular protein opens up new venues in structural characterization, native state recognition, and rational design of proteins.

## 1 Introduction

The native protein structure is uniquely realized because it concurrently satisfies several constraints that are brought about by chain connectivity, the feasibility of non-bonded interactions within the chain and with the solvent, excluded volume effects, and side-chain placements (1,2). Despite this apparent complexity, major structural features of proteins and their functional implications can be studied by coarse-grained models (3–9). Coarse-graining methods have been originally developed to simplify the structure, thereby increasing the attainable length and time scales of computations. Although the underlying principles of coarse-graining of proteins are similar to those used in other materials, i.e., polymers, coarse-graining of proteins are particularly targeted to describe the native state. This has led to the development of various coarse-graining techniques such as Gō models (4,10–13), whose only objective is to fold a given protein assuming a priori knowledge of native topology.

In the current study, we aim to generate coarse-grained model structures comprising assemblies of hard spheres that mimic the geometrical and topological characteristics of real proteins with no reference to specific protein structures. Towards this target, we adopt a construction method with a small number of simple and straightforward rules imposed on a simple cubic (SC) lattice. The ground rules for structure generation are motivated by the average residue packing trends of 210 globular proteins.

The packing characteristics of geometrical objects offer a simple, yet extremely general, paradigm to study condensed matter systems (14–19). Packing approaches have found several uses in protein research. At the most fundamental level, the hydrophobic core’s efficient burial is shown to be a central requirement for protein folding (20–22). Packing density is responsible for the distinct features on the potential energy landscape, and even a simple folded polypeptide chain has an inherently complex and multidimensional landscape, whereas that of a fully extended conformation is featureless (22,23). Tessellation studies showed that residue centers’ space-filling characteristics carry signatures of Bernal packing of hard spheres (24), and free volume distribution resembles that of simple dense liquids (25). At the tertiary structure level, the optimal size of protein domains can be deduced with simple sphere models and efficient burial of the hydrophobic core (26). Finally, proteins have remarkably low intrinsic compressibility, yet they are amenable to unfolding under hydrostatic pressure (27–30). The unfolding pressure is strongly dependent on the free volume distribution, and unfolding is accompanied by a reduction in specific volume (29). For globular proteins, the presence of interior voids are now well documented (31– 33), and more recent analyses of packing hint that the protein core is more open than previously reported (34,35).

In the current study, we investigate the feasibility of using primitive cubic order combined with three sources of disorder: surface/finite size effects, voids, and positional deviations from ideal lattice sites. It is known that the distribution of internal coordinates of C_α_ atoms peaks around preferential values, which are compatible with cubic lattice geometry (36), and crystal lattices are frequently deployed in protein modeling efforts. The underlying lattice provides a basic grid for realizing and updating conformations (10,36–38). There are many folding algorithms based on these ideas (39) that make use of various lattice types. Notably, Covell and Jernigan used a face-centered cubic (FCC) lattice to generate all possible conformations and chose the optimal conformation based on non–bonded pairwise potential energy minimization (40). Similarly, a lattice model based on the diamond cubic lattice (equivalent to an FCC lattice with a two–point basis) was introduced to predict folded conformations at low spatial resolution with no reference to a native state (41).

The presented hard sphere model for structure generation is novel in the sense that it enables us to generate an ensemble of structures at once where each generated member is representative of the structural trends observed in globular proteins. The comparisons rely on the use of experimental coordinates and no attempt is made to elucidate the protein folding mechanisms or related dynamics. Ensemble generation strategy is based on SC lattice clusters with an extra randomly discarded percentage needed to capture the packing trends observed from the experimental coordinates of C_α_ atoms. Excessive introduction of unoccupied sites to the core regions of the generated structures is avoided by two constraints: the single component connectivity of the retained sites and the high surface to volume ratio of the starting clusters. A simple explanation of degeneracy and ruggedness of the potential energy landscape is attempted with these basic tools. Our approach is in a way similar to the Watts–Strogatz approach, wherein one generates small world networks starting from a uniform circular graph and reaches graphs of various clustering, and average path lengths, controlled by the degree of rewiring introduced (42).

To probe the similarities between the generated structures and real proteins, we investigate the distributions of fundamental geometrical and topological descriptors. Through ensemble-based structural comparisons, we are able to quantify the extent of non-superficial similarities between the defect laden cubic order and residue networks. Geometrical comparisons are based on the local bond orientational order (BOO) parameters, and the topological comparisons are based on several network metrics that are directly defined from the moments of the adjacency matrix.

## 2 Computational Methods

The generation of folded protein structures is performed using three simple rules: the presence of a molecular surface, inclusion of free space (voids), and positional deviations from ideal lattice sites. These rules are applied in the order they are listed, and the details of the computational methods used in the generation of the artificial folded protein structures are provided in sections 2.1 In section 2.2, we outline the structural characterization methods used to compare the generated structures to average protein structure.

### 2.1 Generation of Artificial Structures

#### 2.1.1 Formation of Finite Sized Lattice Clusters

In the current study, each lattice point is considered to represent an amino acid (residue) located at the *C*_*α*_ atom. Initially, a finite sized cluster is carved out of the bulk SC crystal with a lattice spacing of 3.8 Å. Throughout this study, 210 different such clusters are randomly generated. These 210 structures are equally selected (35 of each) from 6 different sized clusters: 5×5×6, 6×6×7, 7×7×7, 7×7×8, 8×8×9, and 8×9×9 to capture the size variations in the comparison set of 210 proteins. The shape of the SC clusters is chosen to be approximately cubic because it is the most convenient choice; it is compatible with the basic structure of the underlying SC lattice and it extends equally along the basal directions of the SC lattice. However, it is important to note that clusters of differing geometries were also tried, and that the results do not depend on the shape of the cluster as long as elongated shapes with very high aspect ratios (>1.8) are not used was verified.

#### 2.1.2 Inclusion of Empty Space

Randomly selected residues (coarse-grained beads) are removed from a finite sized cluster until a predetermined void concentration is reached. During the introduction of voids, the structure is checked for connectivity after the removal of each residue. Connectivity is defined such that at any stage during the introduction of voids, a path exists between any two randomly selected residues. Since there are no universal rules regarding the spatial distribution of voids, both internal and surface voids are generated without any constraints. The resulting structures are three-dimensional bead clusters each having a single connected component; these are not necessarily self–avoiding walks because of the way the connectivity is defined during the removal process.

#### 2.1.3 Introduction of Positional Disorder

The structures generated in the previous section are made up of coarse-grained beads located at the lattice points. However, RDFs pertaining to such clusters comprise successions of delta functions and unlike the case for real proteins they do not show any spread. Therefore, it is necessary to relax the residues of the generated structures from their on-lattice positions. This relaxation (the introduction of positional disorder) is achieved through the displacement of coarse-grained beads (residues) via two alternative methods. The first method moves individual beads by targeting the average RDF of real proteins by a reverse Monte Carlo and simulated annealing (RMC–SA) algorithm. The second method invokes displacements by randomly moving residues. The latter is terminated once the average displacement of the beads reaches 0.5 Å. This particular choice ensures that the average displacement is much smaller than the nearest neighbor spacing and is adopted in light of average displacements attained by the RMC-SA method described below.

The reverse Monte Carlo (RMC) procedure is a reconstruction method that aims to evolve a structure until the selected property (or properties) of the evolving structure mimics or best represents the selected experimental property (43,44). Simulated annealing (SA) is a heuristic global optimization method where the probability of accepting solutions is continuously decreased as the solution space is explored. In the current study, the target property of the RMC–SA procedure is chosen to be the average RDF of the 210 basis proteins up to 15 Å distance.

The RMC optimization proceeds with Boltzmann sampling using a simple Metropolis scheme (45). Each attempted move is accepted with a probability (*p*), see Eq. (1), where *ΔE* is the energy difference between the attempted and current configurations of the relaxing structure. The energy function, E, is defined as the integrated squared difference between the RDF of the generated structure, *g*(*r*_*i,t*_), and the average RDF of the 210 basis proteins, 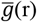, up to 15 Å, see Eq. (2).

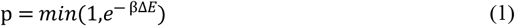

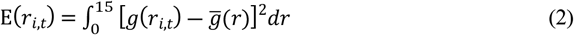

In Eq. (2), the index *i* represents current or attempted structures at step *t*, and the Boltzmann factor is defined as follows:

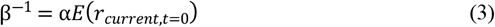

where *α* = 10^−6^ is used as a scaling constant. The simulated annealing is implemented by updating β^−1^ by 0.9 at every 10^3^ steps. The combined RMC–SA procedure is performed for 5×10^5^ steps or until βΔ*E* ≤ 10^−7^ for at least 2×10^3^ consecutive steps.

### 2.2 Characterization of Structural Features

The comparison of the generated structures with an average protein as defined from the 210 globular proteins are performed based on three main features: residue packing, residue bonding anisotropy, and topology (connectivity of residues). The packing consideration is based on the radial distribution function and nearest neighbor considerations. The bonding anisotropy is investigated via the bond orientational order (BOO) parameters. Finally, the contact topology is investigated using network analysis tools. In the next sections, we briefly describe the BOO and the network parameters used in the current study.

#### 2.2.1 Bond Orientational Order

Bond orientational order (BOO) quantifies the anisotropy of bonding; its parameters are defined as the rotationally invariant combinations of bond density expansion coefficients. The bond density expansion is carried out with spherical harmonics which form a basis for the (2*l*+1)-dimensional representation of the rotation group *SO*(3), and therefore, they are well suited to study three-dimensional systems with finite symmetries whose rotational symmetries conform to finite subgroups of the rotational group (46). In the protein community, BOO has been used to define anisotropic statistical potentials, to study the stability of collective motions, and to assign secondary structures (6,47–49).

Originally BOO (50) is defined as a global order parameter, but we will use the local definition that accounts for the bond vectors in the coordination shell for each constituent residue:

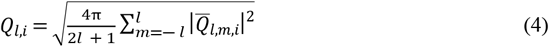

Here 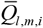 denotes average over all bonds emanating from a residue *i* and is defined as follows:

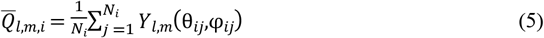

where *Y*_*l,m*_(θ_*ij*_,φ_*ij*_) are spherical harmonics^†^ computed at the polar, θ_*ij*_, and azimuthal, φ_*ij*_, angular coordinates of the unit vector **r**_*ij*_ directed from residue *i* to its neighbor *j*. Note that the final expression of the order parameter *Q*_*l,i*_ in Eq. (4) does not depend on *m* for rotational invariance.

It is possible to systematically define higher order invariants with spherical harmonics (50). The simplest and physically the most interesting higher order invariants are the properly normalized third order invariants given as follows:

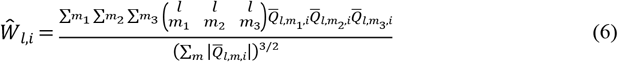

where 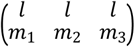 denote the Wigner 3-J symbols and they arise as the maximally symmetric coefficients of three coupled spherical harmonics with vanishing total angular momentum (46). This way the *m* dependency is avoided and the resulting expression is rendered invariant under arbitrary reorientations of the coordinate system. Third order invariants, in their non-normalized form, were first explicitly constructed in the Landau expansions of liquid to nematic transition (51). Later, their normalized form is shown to be central for resolving major symmetries (50). Jarić gives a complete account of the third order invariants in relation to long–range order phenomena within the context of the Landau theory (52).

One remarkable property of these invariants is the persistent sense of parent lattice when computed for crystalline systems. For the three cubic systems; simple cubic (SC), body centered cubic (BCC) and face centered cubic (FCC), all invariants of all *l* coincide up to a sign, whereas hexagonal close packed (HCP) and icosahedral orders are manifested with different shape spectra in *Ŵ*_*l,i*_ (50). Therefore, these parameters are expedient tools to explore the discerning features of the energy landscape. Most notably, the choice of *l* = 6 yields an icosahedral BOO parameter for flat space.

To establish one to one correspondence between two structures, invariants of all orders and of all *l* values have to match (50). The basic difference between different *l* values is the allowable symmetries that can be resolved (50,52). In practice, comparison of the first few degrees is sufficient to see if there is meaningful similarity between two structures. In the current work, we use the second order invariants of *Q*_*l,i*_ and the third order invariants *Ŵ*_*l,i*_ with *l* = 2, 4, and 6.

Steinhardt et al. offered BOO as a means for “shape spectroscopy”, and conventionally, BOO parameters constructed with *l* = 6 were used in the literature because it is the lowest *l* that can distinguish cubic and hexagonal configurations as well as the non–crystallographic icosahedral symmetry. Whether a given *l* representation can distinguish a given symmetry follows from the isotropy groups of O(3) (52). Isotropy group comprises the elements of O(3) that leaves the (2*l*+1)– dimensional vector [*Q*_*l*, − *l,i*_,*Q*_*l*, − *l* +1,*i*_,…, *Q*_*l,l,i*_] invariant. If no such group exists for a given *l* the corresponding order parameters *Q*_*l,i*_ and *Ŵ*_*l,i*_ will be identically zero.

#### 2.2.2 Network Analysis

Network view provides a convenient framework for comparing two systems in terms of inter–unit interactions. In the current study, networks are constructed from the residues of the generated structures by connecting residues that were within a 7 Å cut–off distance of each other. The choice of cut-off distance is motivated in part because 7 Å allows non–bonded interactions within the two nearest coordination shells to be included (53–55). This cut-off value is also consistent with previous Delaunay tessellation and cut-off scanning studies (56,57).

The contact topology of a residue network is stored in the *N*×*N* adjacency matrix ***A***, where the *A*_*i,j*_ term is either equal to one or zero according to the presence or absence of interactions between the *i*^*th*^ and *j*^*th*^ residues, respectively (54). Analysis of the adjacency matrix and its variants (such as the Laplacian or the normalized Laplacian) leads to a wealth of information. For example, a statistical mechanical treatment yields information on residue auto- and cross-correlations (58), and the neighborhood structure of a residue and how information propagates to further neighbors (55). Network models were also used to identify adaptive mechanisms in response to perturbations (59), predict collective domain motions, hot spots, and conserved sites (58,60–62).

In terms of network characteristics, proteins share common structural similarities with other self-organizing condensed matter systems (53). A quantity termed “relative contact order”, which may be derived from the adjacency matrix, is shown to highly correlate with the folding rates measured for many two-state folders (63).

In the current study, distributions and averages of local network parameters such as degree (*k*_*i*_, number of neighbors of residue *i*, Eq. (7)), the nearest neighbor degree (*k*_*nn,i*_, the number of neighbors of the neighbors of residue *i*, Eq. (8)), and the clustering coefficient (*C*_*i*_) of residue *i*, Eq. (9), were monitored. In addition, a global parameter, the shortest path length (*L*_*i*_) of residue *i* is also monitored (Eq. (10)). These network parameters are defined as follows:

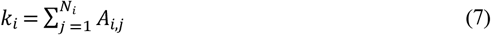

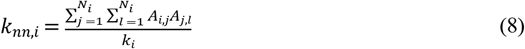

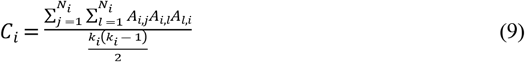

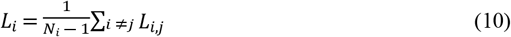

For proteins and other self–assembled systems, the nearest neighbor degree (*k*_*nn*_) is an important classifier and is a linearly increasing function of degree (*k*) with a slope that is equal to the average clustering coefficient 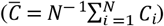 (23). The clustering coefficient (*C*_*i*_) is indicative of the amount of local structure present in the immediate neighborhood of a residue. The clustering coefficient has been shown to converge to a value of approximately 1/3 for residues that are buried in the protein (54). In Eq. (10), *L*_*i,j*_ is the minimum number of steps between residues *i* and *j*. The average shortest path length 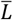 calculated from *L*_*i*_ values is a global parameter displaying the efficiency with which different parts of a protein communicate with each other and is shown to highly correlate with residue fluctuations (54).

The Laplacian (***L***) is used extensively in the graph theory literature (64,65) and is defined as follows:

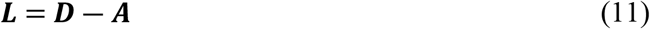

where ***D*** is a diagonal matrix with *D*_*i,i*_ = *k*_*i*_. The Laplacian is a positive–semidefinite matrix and its eigenvalue spectrum is routinely used for quantitative characterization of networks. In the current study, the normalized Laplacian (***L***^*^) is used.

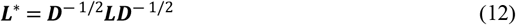

The main utility of the normalized Laplacian comes from the fact that all of its eigenvalues exist within the [0–2] interval (66,67). Therefore, the spectra of networks of differing sizes can be directly compared if the Laplacian matrix is normalized. More specifically, the presence of an eigenvalue with *λ*= 2 signals that the graph has a bipartite connected component. The presence of an eigenvalue with λ = 1 shows that there are two vertices with identical connections (vertex doubling); and more generally, the multiplicity of eigenvalues with λ = 1 quantifies the extent of replication among structural motifs. The number of eigenvalues with λ = 0 is equal to the number of connected components. Finally, the smallest no-zero eigenvalue can be used as a quantitative measure of the difficulty to disconnect the network (the greater the value of the first non–zero eigenvalue is, the harder to disconnect the network becomes), and therefore, is a measure of collectivity in the network. Here, a connected component refers to a set of nodes for which there is at least one path among any pair of nodes in the set (65).

## 3 Results and Discussion

Throughout this study, we will show that at least three simple requirements are necessary to generate an artificial structure (from a crystalline template) that mimics the behavior of average folded protein:

1. The presence of a molecular surface (finite size effect)
2. The inclusion of free space
3. Positional disorder

Furthermore, it will be shown that although the third requirement is important in reproducing the average protein radial distribution function, it has almost no impact on a variety of other structural features observed in real folded proteins.

As mentioned earlier, instead of targeting a specific protein to mimic (or identifying) its structure, we target the average behavior of many proteins based on the experimentally determined coordinates. Therefore, as a first step, it is necessary to define a set of proteins (“the basis set”) from which the average structural features of real proteins can be defined and the generated structures can be compared against.

### 3.1 The Average Protein Structure and the Selection of Lattice Template

To define an average protein structure, a set of 210 globular proteins was selected from 595 proteins, which are representative of four major protein folds with less than 25% sequence homology among them (54,68). The selection of the basis proteins was decided according to their globularity, which is investigated by a principle component analysis of amino acid C_α_ coordinates of these 595 proteins via singular value decomposition (69). The proteins, whose ratio of the largest and lowest singular vectors (σ_*max*_ /σ_*min*_) is less than 1.8 were selected into the basis protein set. The ratio of 1.8 is based on hen egg white lysozyme (PDB Code: 193L), which is one of the most studied proteins. This selection criterion provided 210 proteins after eliminating rod–like proteins from consideration. The details of the basis set proteins are provided in the Supplementary Data section.

We first compare defect free crystalline systems using BOO to see whether the local coordination characteristics in proteins can be directly reproduced by any of them. The average BOO parameters, 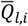 and 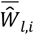, that were calculated from the 210 basis protein structures are shown in Figure 1. The average 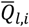 and 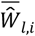 values suggests that the average residue neighborhood is not a direct analogue of crystals (50). Icosahedral order is also deemed to be absent for similar reasons because perfect icosahedral order requires that 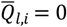 for *l* = 2, 4, and this is not the case as can be seen in Figure 1. Since comparisons with ideal crystalline lattices fail, one should consider drawing comparisons to systems that contain defects such as point defects (voids) or positional disorder.

**Figure 1.**
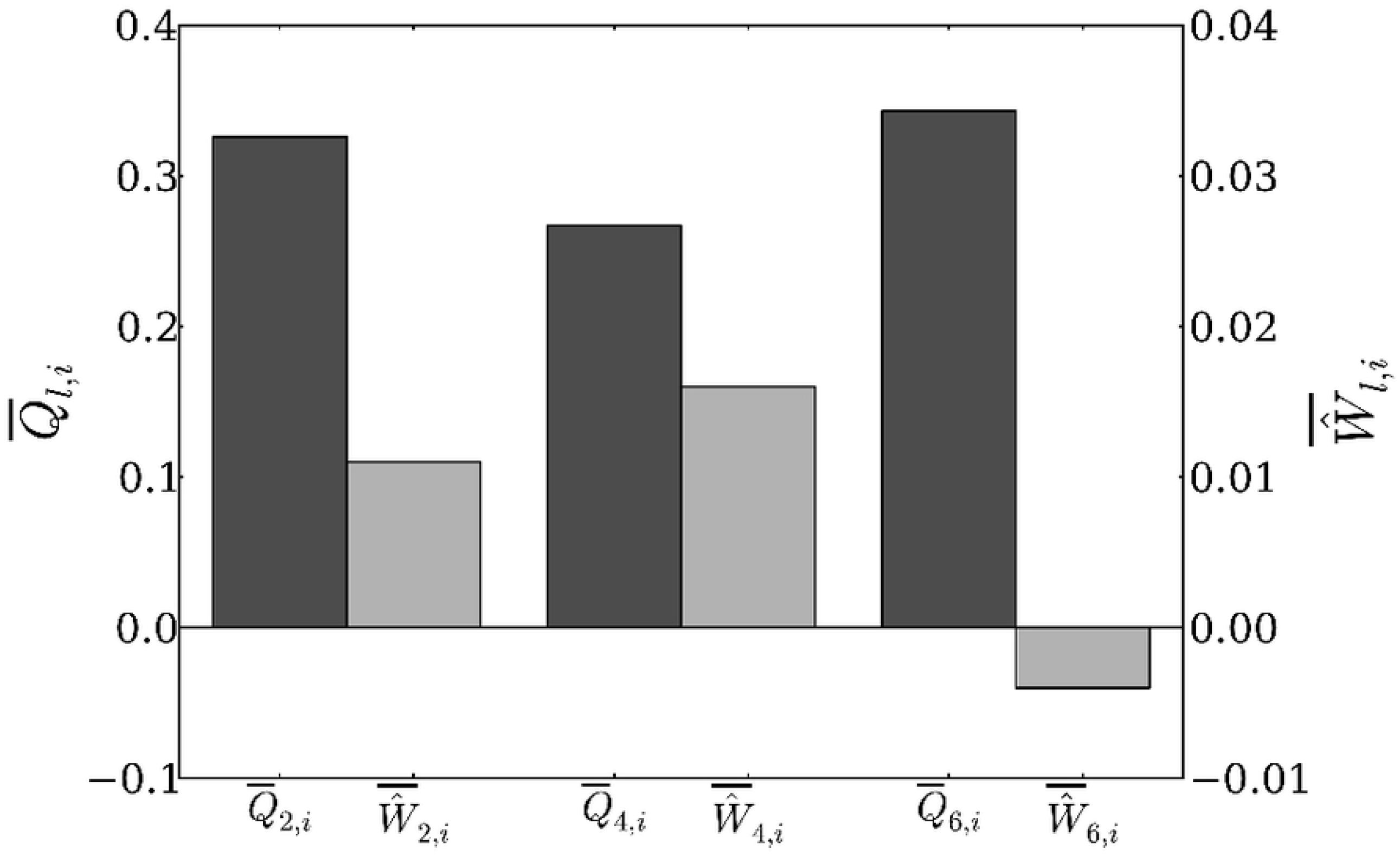
The average BOO parameters, (dark grey) 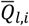 and 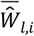 (light grey), for *l* = 2, 4, and 6 for the 210 proteins used as the basis set.

The RDF is an important metric of any packing consideration; therefore, any attempt to involve packing arguments should naturally start by an investigation of the RDF. The RDF of proteins (coarse-grained at the amino acid level) is known to have generic characteristics (70,71). The individual RDFs (gray shaded region) and the average RDF (solid line) of the 210 basis proteins are shown in Figure 2.

**Figure 2.**
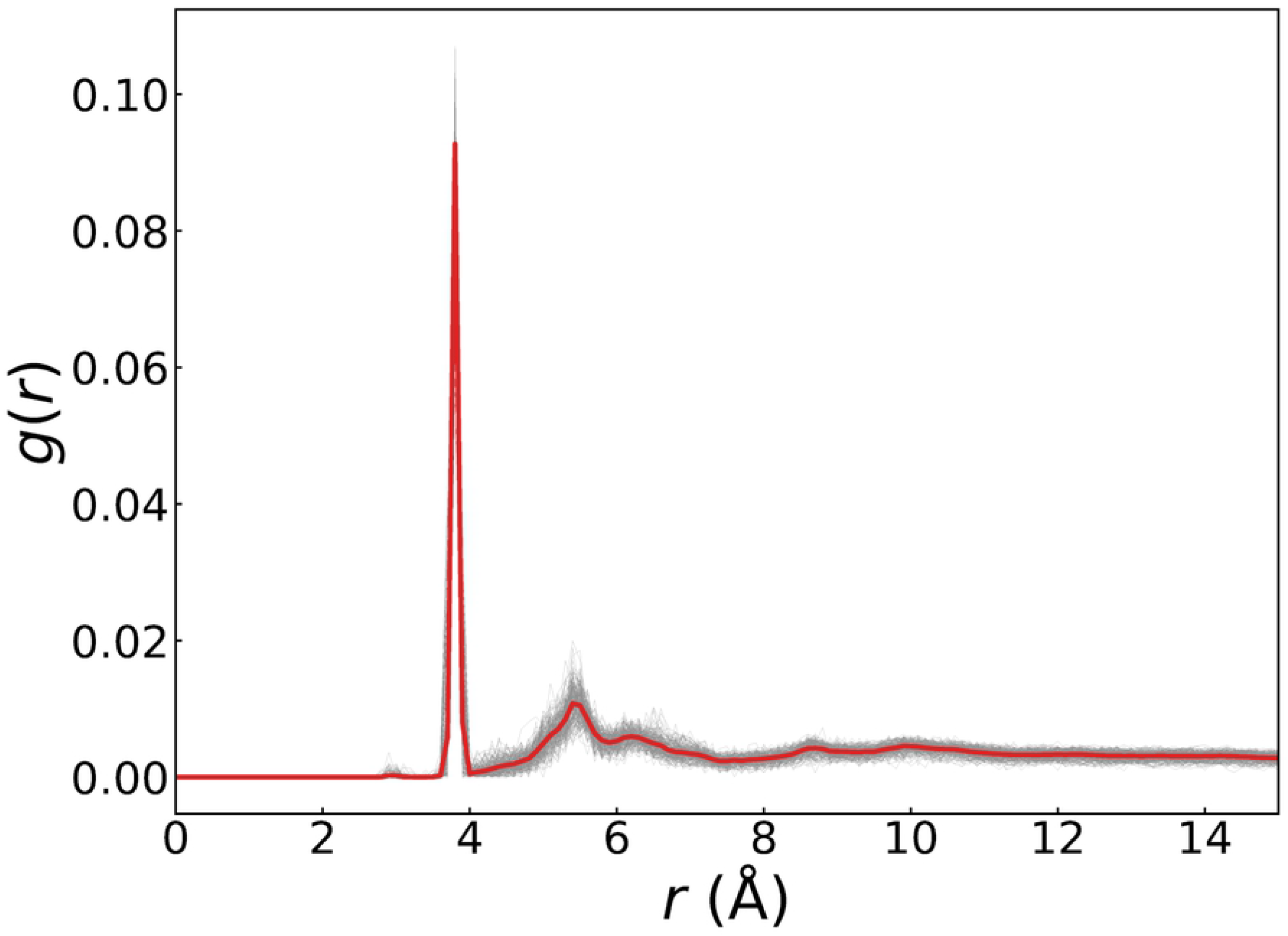
Radial distribution functions, *g*(*r*), of 210 globular proteins (thin gray lines) and their average radial distribution function (bold line) calculated with a bin size of 0.1 Å.

The average RDF is marked by a sharp first coordination peak at 3.8 Å and is the due to the bonds between nearest neighbors on the main chain. In addition, a less prominent secondary peak is observed at ∼5.6 Å. The ratio of the first and second peak locations puts fundamental constraints on the placement of neighboring residues on a lattice. Fortunately, for the FCC and SC lattices, the ratio of the first and second nearest neighbor distances is equal to 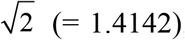, which is quite close to the ratio obtained from the average protein RDF (5.6/3.8 = 1.4737). Therefore, it is plausible to accept FCC and SC lattices as candidate template lattices. However, when the average number of nearest and next-to-nearest neighbors of nodes in proteins is calculated from the area underneath each peak, it becomes evident that the first coordination shell needs to be less crowded than the second coordination shell. However, in FCC, which is a close packed structure, the opposite is true and therefore, the FCC lattice cannot be considered as a template lattice. This leaves the SC lattice as the only candidate. Although this selection might be viewed as counterintuitive, it has been shown that the bending and torsional angles of proteins coarse-grained at the amino acid level show peaks at highly preferential values agreeing with the geometry of the SC lattice (36). Therefore, the selection of SC lattice as the template for the generation of coarse-grained, artificial protein structures is not a coincidence and it is based on multiple structural considerations of real proteins.

### 3.2 Generation of Finite Sized Clusters

The first step in the generation of artificial proteins is the carving of finite–sized clusters from the SC lattice template. This step is required to capture the finite size of proteins. However, it is clear that any finite–sized SC cluster that is occupied with residues at each lattice point would be too dense compared to proteins. The cut-off variation of the average contact numbers (degree, *k*) for the 210 basis proteins and various SC clusters as a function of occupancy are shown in Figure 3, which clearly shows that the perfect SC cluster at 100% occupancy is too dense compared to the average behavior of real proteins. Interestingly, at 40% occupancy (after randomly removing 60% of the lattice points), the average contact density of 30 SC clusters shows almost the same cut-off variation as that of the average protein.

**Figure 3.**
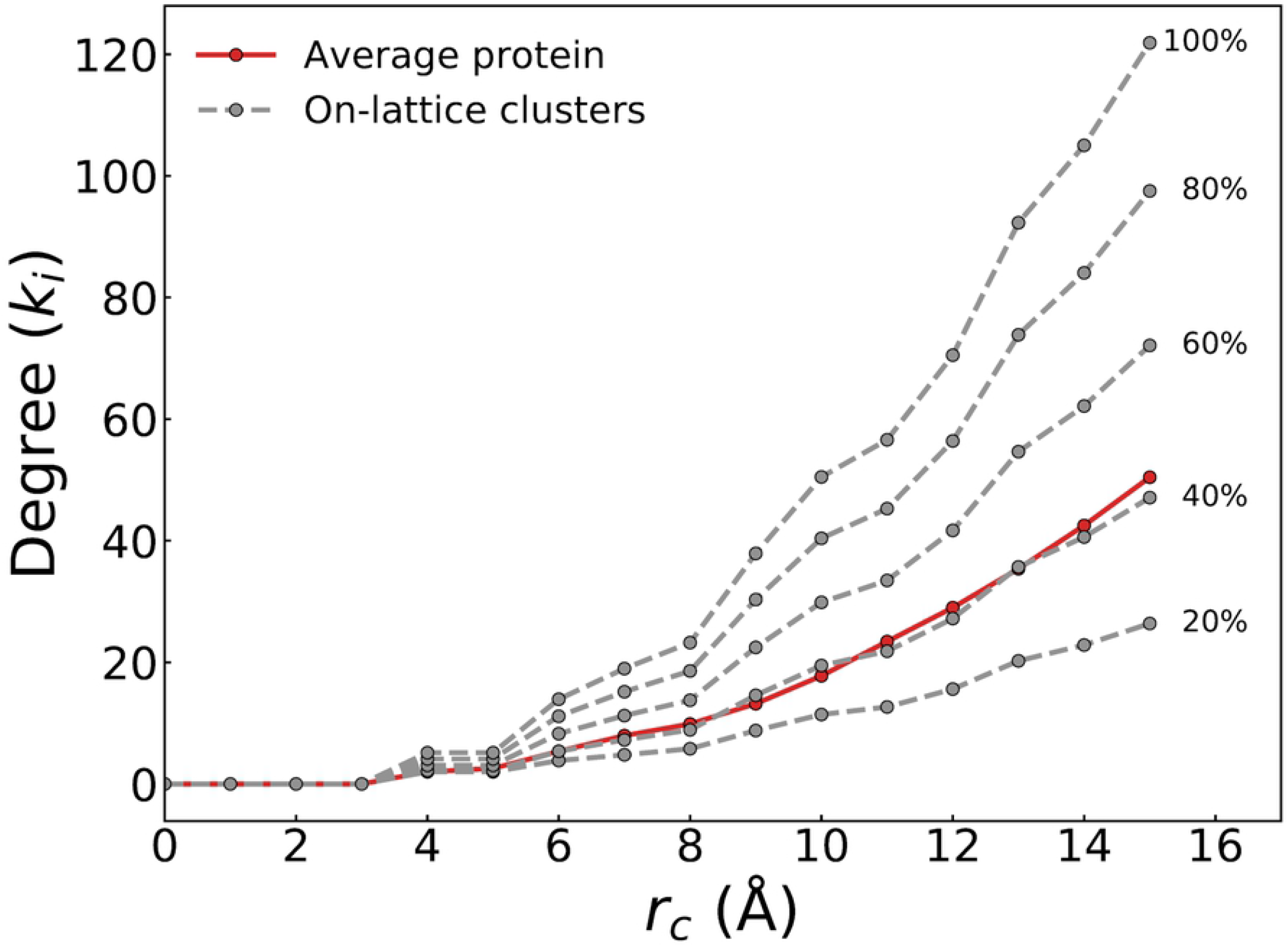
The dependence of the average degree (*k*) on cut–off distance for 210 basis proteins (solid line) and SC clusters (dashed lines) at various occupancies.

This finding suggests that in order to create structures similar to proteins, the SC clusters should be emptied down to ∼40% occupancy. We also note that 40% occupancy as well as the 60% void concentration are both above the percolation threshold of 25% of the SC lattice (72). This suggests that once the SC cluster is emptied down to 40% occupancy, the remaining residues could form a connected network of points, which is an absolute requirement for any protein–like structure to be considered viable. However, random removal of lattice sites does not guarantee that the remaining lattice sites form a connected network; therefore, an additional constraint is imposed during the random removal procedure to maintain the connectedness of the remaining lattice points. In the current study, the connectedness is defined such that there must be at least one path between any two remaining beads on the cluster through nearest neighbor contacts. Finally, three-dimensional networks are constructed by introducing additional bonds among the constituent beads that account for the second coordination shell. Construction process is illustrated on a 7×7×7 parent cluster in Figure 4.

**Figure 4.**
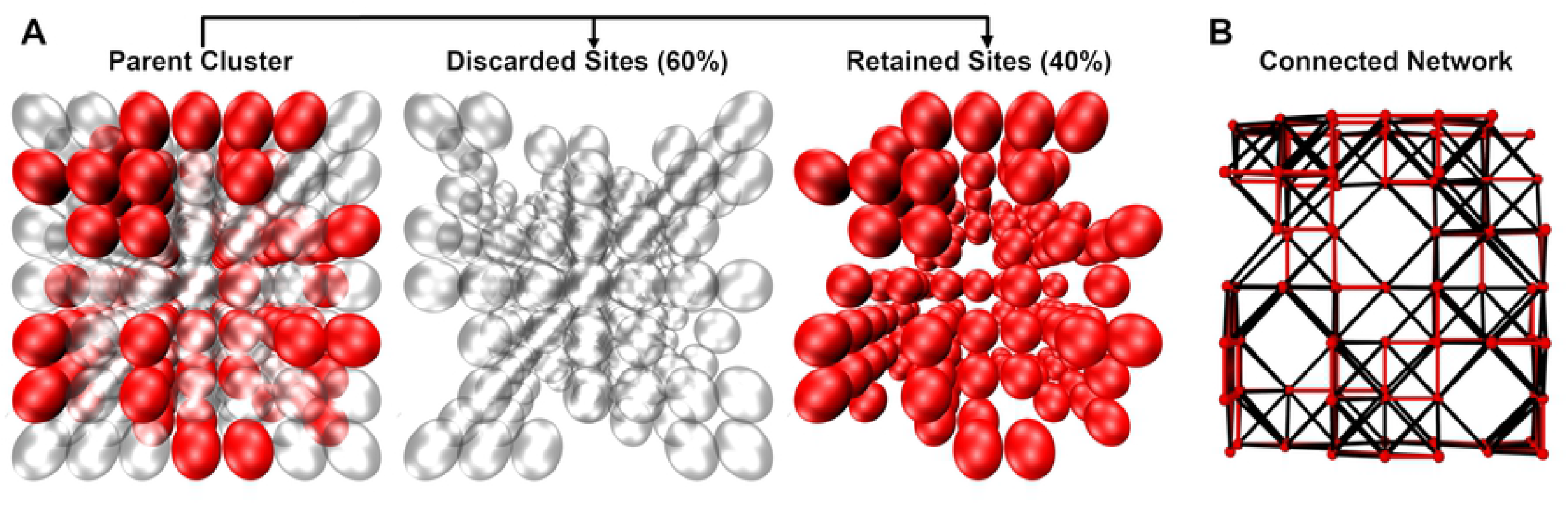
Construction of a 3d artificial network on a parent 7×7×7 SC cluster. (a) Parent cluster is emptied down to 40% occupancy by discarding 60% of the initial sites. The remaining structure comprises 138 beads which form a single connected component through nearest neighbor contacts. (b) Additional contacts are introduced by adopting a 7 Å cut-off. Nearest neighbor contacts are colored with red whereas the rest are shown in black.

Since the residues of the structures generated on the SC lattice template are located on regularly spaced lattice positions, their RDF would show up as a succession of delta functions with no spread irrespective of occupancy, see Figure 5(a). It is well known that positional disorder (displacement of lattice points from their ideal locations) creates spread in RDF peaks. Therefore, the random displacement and the RMC–SA procedures are used to evolve the (emptied) clusters.

**Figure 5.**
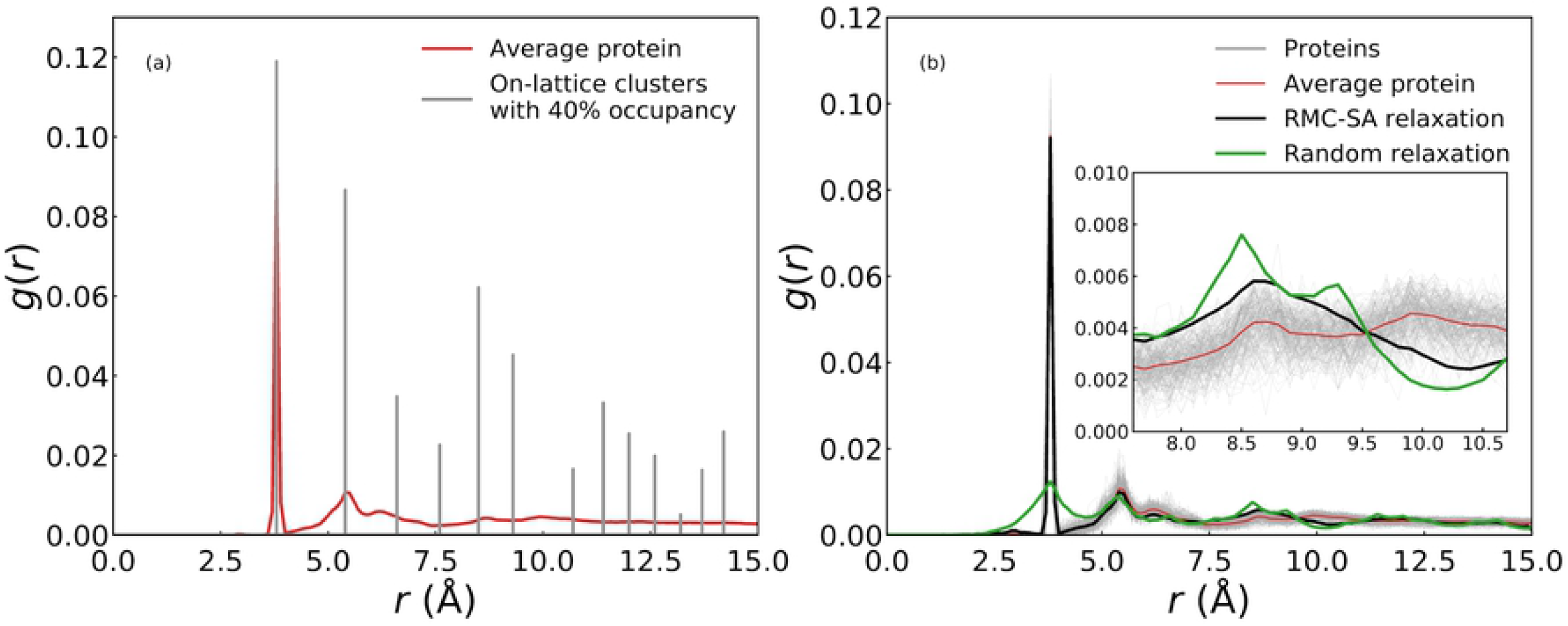
(a) Radial distribution functions, *g*(*r*), of the average protein and an example on–lattice cluster structure at 40% occupancy before introducing positional disorder. (b) Comparison of the average radial distribution functions of 210 basis proteins (circles) and 210 clusters at 40% occupancy after relaxation of the on–lattice residue locations with random displacements (dashed line) or RMC–SA procedure (solid line) calculated using a bin size of 0.1 Å. Inset shows the discrepancy between real proteins and generated structures.

### 3.3 Introduction of Positional Disorder

The comparison of the average RDF for 210 basis proteins and 30 clusters at 40% occupancy relaxed with random displacements or with RMC–SA procedure is shown in Figure 5(b). The relaxation procedure performed with random displacement of residues does not reproduce the average RDF of the basis proteins, the main difference between the two RDFs being at the first coordination peak. On the other hand, because the RMC–SA relaxation procedure uses the average protein RDF as its target function, the RDF of the cluster relaxed with RMC–SA procedure successfully matches that of the average protein. The main discrepancy between the two RDFs is around the distance range of 7.6–10.7 Å (see inset to Figure 5(b)). The sub-range from 7.6 Å to 9.3 Å corresponds to three on-lattice peaks that are representative of the 4^th^, 5^th^ and 6^th^ closest neighbors on the SC lattice (Figure 5(a)), and it is clear that the RMC–SA relaxation procedure is not able to reduce the high number of initial contacts that were present in the on-lattice structure within this distance range. The reasons for this outcome are not known. However, one possible explanation might be that the connectedness constraint imposed during the introduction of voids is not strict enough to produce a linear chain, and therefore, side branches materialize after the introduction of voids. Because the relaxation procedure does not remove branches, their existence leads to a more compact structure and a slightly higher density within the core of the cluster. From 9.3 Å to 10.7 Å, the source of discrepancy is more straightforward. This range is the empty region between the 6^th^ and 7^th^ nearest neighbors on SC lattice (see Figure 5(a)) and it is larger than the average displacements introduced with the two relaxation methods. Therefore, RDF functions pertaining to the generated structures attain lower values in comparison to the proteins.

The radial displacements of all residues and the average radial displacement as a function of distance from the center of mass are shown in Figure 6 for the two relaxation procedures employed. Neither the individual nor the average radial displacements are ever greater than the average nearest neighbor distance (3.8 Å) between residues regardless of the location of the residue from the center of mass (buried at the protein core or at the free surface). However, the residues at the free surface experience (on average) slightly greater displacements compared to those closer to the center of mass during the RMC–SA relaxation, see Figure 6(a). When the relaxation is performed with completely random displacements, the difference between the core and surface disappears, see Figure 6(b). Although the RMC–SA relaxation procedure do not inherently impose any location specific constraints, it is clear that the unintended consequence of trying to fit the average RDF of the basis proteins (which is the target function of the RMC–SA procedure) leads to this outcome. This behavior is also compatible with the extra mobility that would be enjoyed by residues residing on the surface of real proteins.

**Figure 6.**
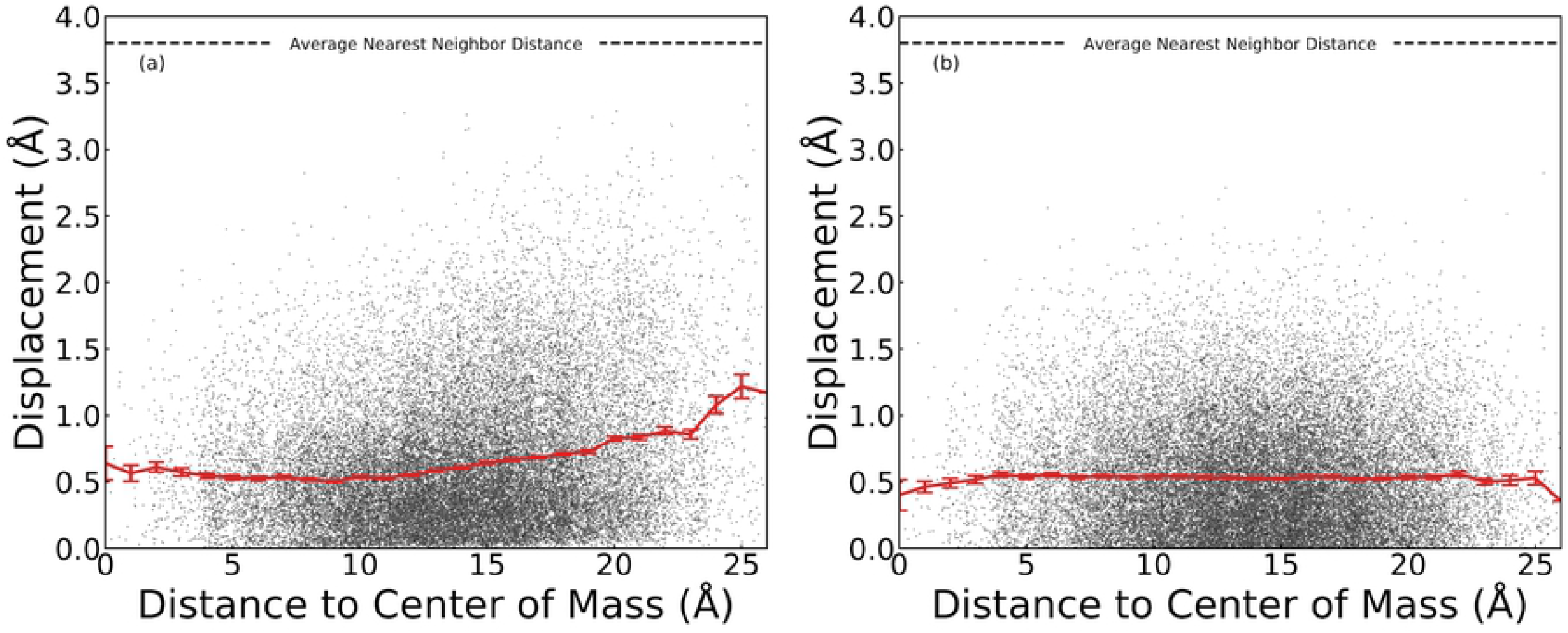
Distribution of the radial displacements of all simulated residues as a function of radial distance from the center of mass of the cluster during (a) RMC-SA and (b) random displacement procedure. Solid curves are moving averages and the error bars represent the standard error.

### 3.4 Comparison of Structural Features

#### 3.4.1 Bond Orientational Order Analysis

RDF is a reduced (directionally averaged) pair distribution function. Although it can distinguish different condensed phases of matter such as crystals, liquids, and gases, a multitude of structures could realize the same RDF. Therefore, an optimization which only takes the RDF as its objective function, is not guaranteed to generate a unique structure. To understand if the generated structures are comparable to real proteins, one needs to match other properties. For this reason, additional comparisons are performed both geometrically and topologically. The geometric comparisons are based on BOO and the relevant order metrics 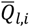 and 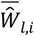 for *l* = 2, 4, and 6. BOO parameters quantify the differences in anisotropy of the local environment. All BOO parameters are computed with a cut-off distance of 7 Å. This choice is based on previous Voronoi studies which showed that a nearest neighbor definition should not be limited to the intra-chain neighbors (that are separated by 3.8 Å) only and that all neighbors within 7 Å should be considered (24,56). In addition, this choice of cut-off distance accounts for nonbonded interactions within the first two neighbor shells. The results of the BOO analysis are shown in Figure 7 and Table 1. There is nontrivial correspondence between BOO parameters computed for all degrees of *l*. For each *l* studied, on–lattice and relaxed clusters produced distributions that are similar to that of the average protein in terms of shape (unimodal distributions) and range/variation. In general, the real proteins produced *Q*_*l,i*_ distributions with stronger peaks. For *l* = 2 and 4, the average protein has slightly lower mean values for *Q*_*l,i*_. This is most likely a manifestation of an underlying chain, which puts restrictions on the organization of nonbonded contacts. Additionally, for *Q*_2,*i*_ parameter, the fact that it cannot resolve cubic symmetry might contribute to the observed discrepancy. Finally, the BOO parameters have a size component which scales inversely with the contact number, and because proteins have a slightly higher average contact number, the protein distributions might translate to slightly lower average BOO parameters although this is not observed for *Q*_*l,i*_ for *l* = 6. In general the correspondence between the average protein and the relaxed structures is better for *Ŵ*_*l,i*_ than it is for *Q*_*l,i*_ for *l* = 4 and 6. The (unrelaxed) on-lattice clusters show a strong peak at *Ŵ*_2,*i*_ = 0 (Figure 7(d)) because *Ŵ*_2,*i*_ vanishes for *l* = 2 in a cubic cluster (50) and the starting structure will have points that sit at an ideal cubic environment. Although this peak quickly disappears once the structure is relaxed through random moves and RMC–SA algorithm, the relaxed probability distribution functions show the opposite variation in *Ŵ*_2,*i*_ than that is observed for the average protein structure. This discrepancy seems to be the most prominent disagreement between the proteins and relaxed structures.

**Table 1.**
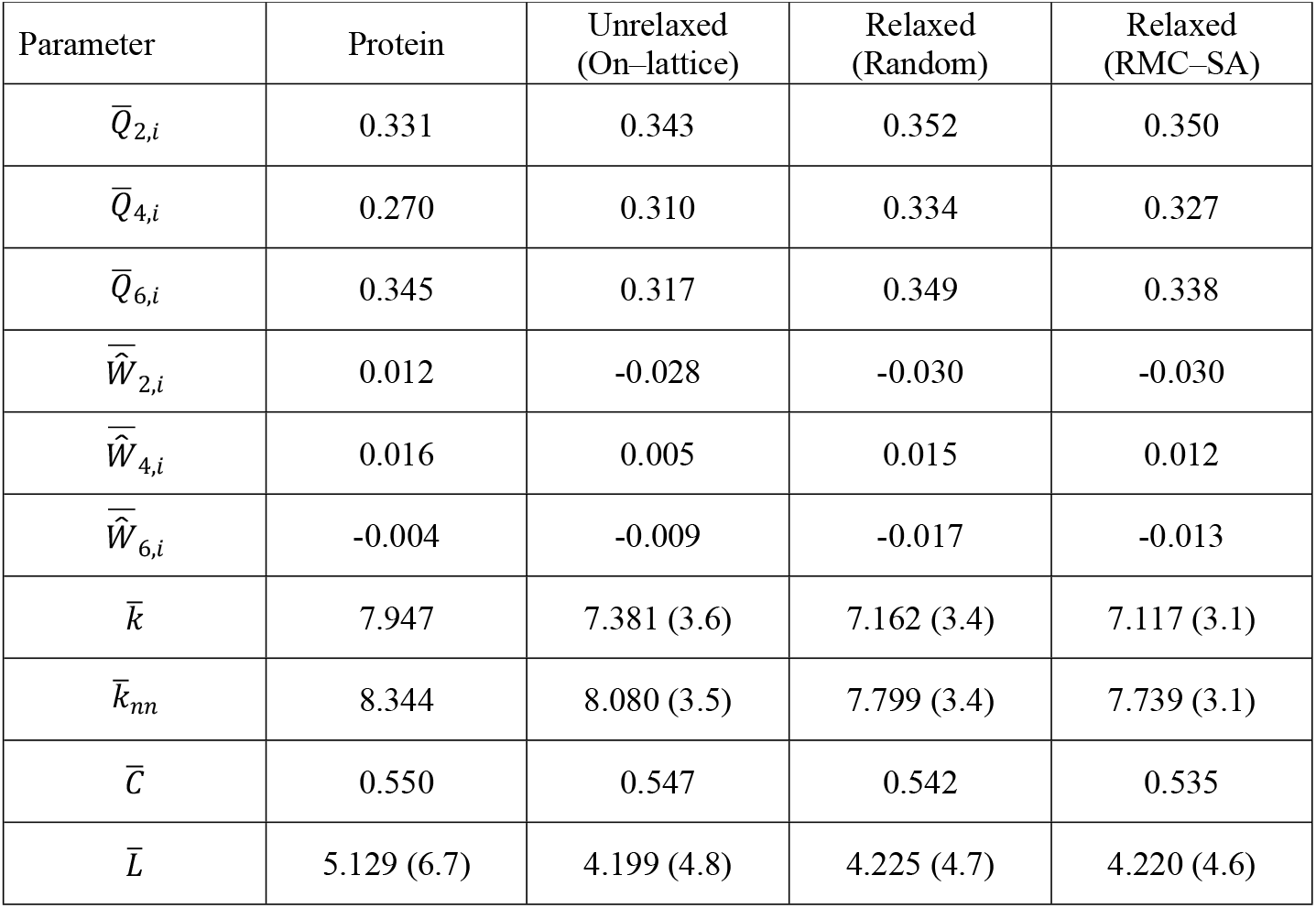
Mean values of various Bond Orientational Order and network parameters for 210 basis proteins, unrelaxed on-lattice clusters, clusters relaxed with random move algorithm, and clusters relaxed with RMC–SA algorithm. All generated structures have 40% occupancy. The average eigenvalue of the normalized Laplacian is not shown because it is always equal to one by definition. Standard error of the last digit is shown in parentheses only when it is greater than 1%.

**Figure 7.**
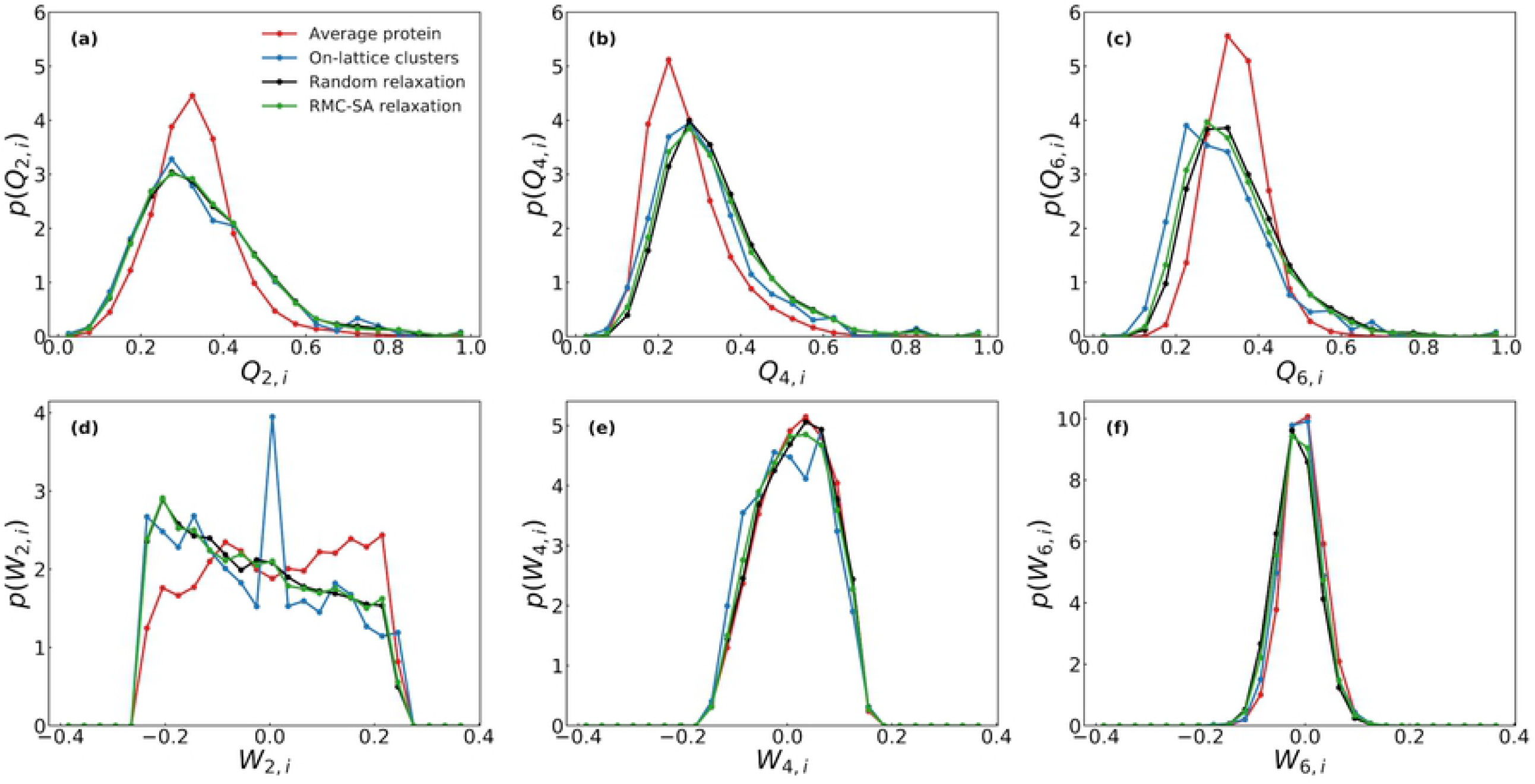
The comparison of various BOO parameter distributions of average protein and various generated structures with 40% occupancy for *l* =2, 4, and 6.

It is clear that although RMC–SA algorithm greatly improves the agreement between the generated structures and real proteins in terms of the RDF, it has much smaller effect on the BOO parameters. Relaxation with random moves gives almost exactly the same BOO parameter distributions as those obtained with RMC–SA algorithm. In fact, with the exception of the sharp peak observed at *Ŵ*_2,*i*_ = 0, the on–lattice structures show distributions quite similar to those of the relaxed structures. This is an important finding in the sense that the template lattice (at 40% occupancy) seems to provide most of the necessary structural similarities. Although structurally neither the on–lattice nor the relaxed structures are exactly the same as the average protein structure, the similarities are remarkable. We should also note that the generated structures could be improved if more rigorous optimization methods are employed. For example, the RMC–SA algorithm could be made to optimize against a collection of structural properties in addition to RDF, and as a result, better agreement could be obtained between the structural features compared. However, because the current study aims to identify the simplest set of rules to generate structures similar to folded proteins, a more rigorous approach is not pursued at this time.

To quantify the extent of similarity between the generated structures and the average protein structure, two–sample Kolmogorov–Smirnov tests are performed. The Kolmogorov– Smirnov test is a nonparametric statistical test that compares two independent distributions and is sensitive to location and shape of distributions. The Kolmogorov–Smirnov test calculates the maximum distance (*D*) between two cumulative distribution functions (CDFs). In our case, we compared the generated structures to real proteins; therefore, *D*_*x*_ is calculated for various properties (*x* stands for various BOO or network parameters investigated) between the average cumulative distribution functions of 210 proteins and 30 generated structures as follows (73):

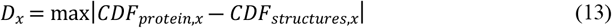

In the two–sample Kolmogorov–Smirnov test, the null hypothesis is rejected with a confidence level (*α*) if the following inequality holds:

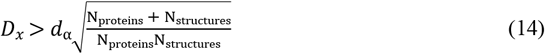

where *N*_*proteins*_ and *N*_*structucres*_ are the number of data points making up the respective CDFs. The value of *d*_*α*_ (tabulated in (73)) depends on *α* (a value of 0.05 is used generally) and sample size. In the current study, the null hypothesis is that both groups were sampled from populations with identical distributions; therefore, we would like to see that this null hypothesis is not rejected. To provide a confidence level to (accepting or rejecting) the null hypothesis, we calculated *p*-values, which indicate the probability of obtaining a result equal to or more extreme than what is actually observed assuming that the null hypothesis is true. In general, the null hypothesis is rejected if the *p*-value is less than a predetermined confidence level (generally 0.05 or 5%). The results of the two-sample Kolmogorov–Smirnov tests are shown in Table 2. It is clearly seen from the *p*-values that the null hypothesis (that the two samples, proteins vs. various generated structures, are drawn from identical distributions) is not rejected for any of the generated structures at 40% occupancy. As a result, it can be stated that the correspondence obtained between proteins and various generated structures is a statistically significant observation.

**Table 2.** Two–sample Kolmogorov–Smirnov test parameters comparing various structural property distribution functions of proteins with those of unrelaxed (on–lattice) and relaxed clusters. All generated structures have 40% occupancy.

| Parameter | Unrelaxed<br>(On-lattice) |  | Random Relaxation |  | RMC-SA Relaxation |  |
| --- | --- | --- | --- | --- | --- | --- |
| | $D$ | $p$-value | $D$ | $p$-value | $D$ | $p$-value |
| $Q_{2,i}$ | 0.113 | 0.999 | 0.123 | 0.996 | 0.117 | 0.998 |
| $Q_{4,i}$ | 0.155 | 0.954 | 0.242 | 0.539 | 0.215 | 0.691 |
| $Q_{6,i}$ | 0.249 | 0.501 | 0.116 | 0.998 | 0.161 | 0.938 |
| $\hat{W}_{2,i}$ | 0.140 | 0.945 | 0.123 | 0.983 | 0.121 | 0.986 |
| $\hat{W}_{4,i}$ | 0.073 | 1.000 | 0.011 | 1.000 | 0.028 | 1.000 |
| $\hat{W}_{6,i}$ | 0.054 | 1.000 | 0.134 | 0.961 | 0.097 | 0.999 |
| $k_i$ | 0.139 | 0.974 | 0.154 | 0.937 | 0.158 | 0.924 |
| $k_{nn,i}$ | 0.117 | 0.997 | 0.176 | 0.848 | 0.189 | 0.783 |
| $C_i$ | 0.145 | 1.000 | 0.127 | 1.000 | 0.132 | 1.000 |
| $L_i$ | 0.283 | 0.291 | 0.276 | 0.319 | 0.277 | 0.315 |
| $\lambda$ | 0.047 | 1.000 | 0.049 | 1.000 | 0.053 | 1.000 |

#### 3.4.2 Network Analysis

The probability distributions of various network parameters of the average protein and various generated structures are presented in Figure 8. Comparison of the degree (*k*_*i*_) distributions suggests that the proteins have slightly greater contact number within the 7 Å cut-off distance, Figure 8(a). This observation may be attributed to the fact that the coarse-grained protein is a self-avoiding chain, whereas, the connectivity criteria used during the generation of the clusters do not impose a self–avoiding walk constraint. The effects of the connectivity criteria are also seen in the nearest neighbor degree (*k*_*nn*_), clustering coefficient (*C*), and shortest path length (*L)* distributions (Figure 8(b), (c), and (d), respectively). However, despite the slight differences in the local and global network parameters, the generated clusters capture the network specific properties of proteins considerably well.

**Figure 8.**
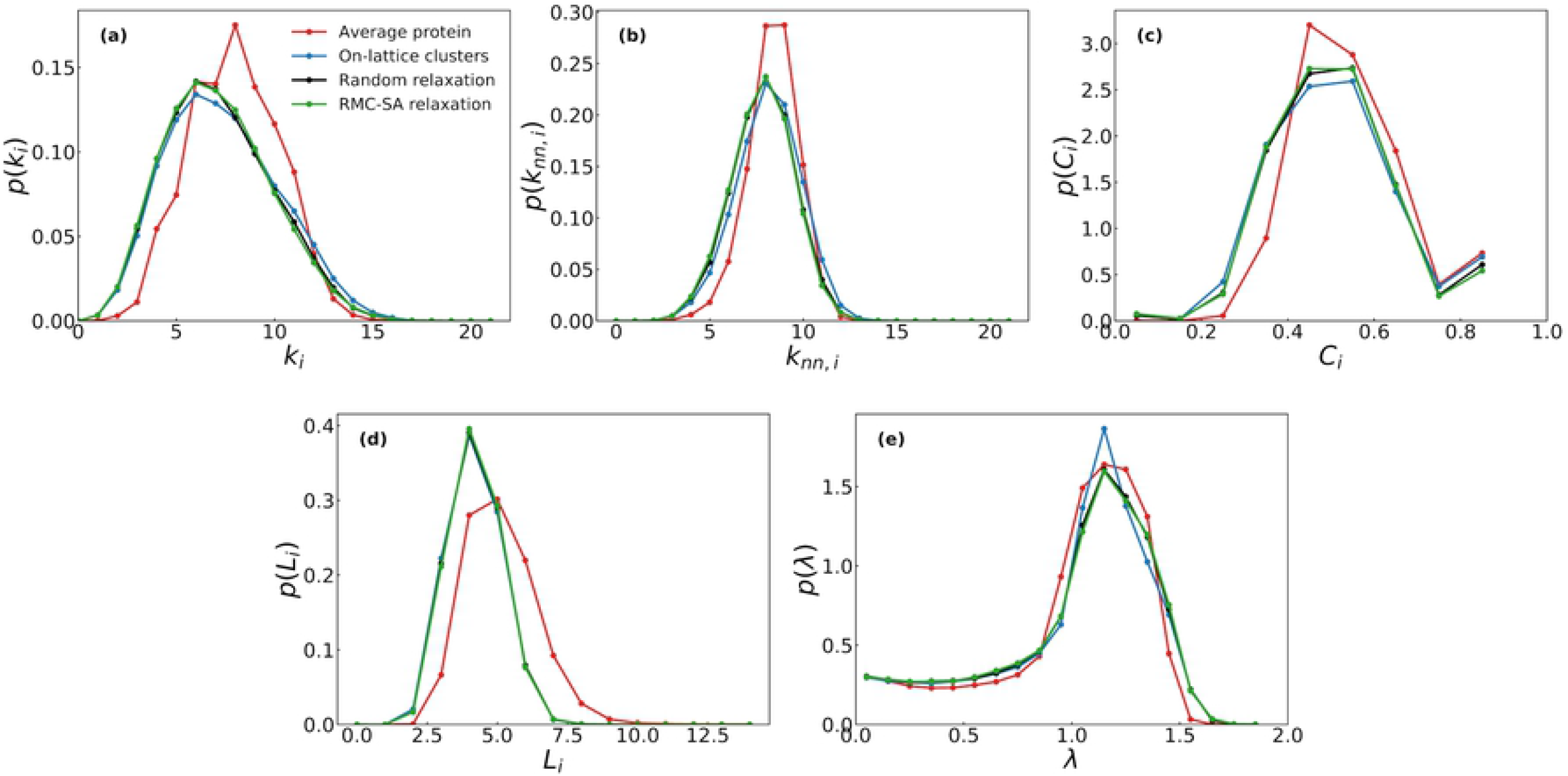
Comparison of the distributions of (a) the degree (contact number), (b) the nearest neighbor degree, (c) the clustering coefficient, (d) the shortest path length, and (e) the eigenvalues of the normalized Laplacian for average protein and generated clusters with 40% occupancy.

Figure 8(e) displays the eigenvalue distribution of the normalized Laplacian for the generated clusters and the average protein. The eigenvalue distribution of the average protein is characterized by a wide peak in the region 1.0 < *λ* < 1.5, overlaid by a long tail at 0 < *λ* < 1.0. The eigenvalues in the proximity of *λ* = 0 carry signatures of collectivity in the system and indicates the ease with which the network may be disconnected; the generated structures capture this region remarkably well. In addition, the region where *λ* > 1 is marked by a wealth of local motifs that are realized by secondary structural elements, and therefore, it is expected to cause the biggest discrepancies. However, the generated clusters successfully mimic the behavior of the average protein structure in the region *λ* > 1. Finally, the overall distribution of the generated clusters is observed to be typical of the molecular structure networks (53,74).

The results of the Kolmogorov–Smirnov tests on various network parameters are also listed in Table 2. Based on the test results, the null hypothesis is once again not rejected, and therefore, the possibility that the network parameter distributions of the real proteins and various generated structures come from the same distribution cannot be overruled.

### 3.5 Influence of Occupancy

The results of the structural comparison of the unrelaxed (on–lattice) and relaxed generated structures to average of 210 real proteins reveal that with the exception of a few differences, the relaxation of the residues from their on–lattice locations do not result in substantial differences in the BOO and network properties. Thus, most of the structural features are inherited from the lattice template used to create these generated clusters. One of the rules imposed on the generation procedure is to empty the clusters such that the occupancy would be 40%. This leads to an excellent agreement between the average degree (contact number) of the on–lattice clusters and real proteins (see Figure 3). However, it is important to understand the effect of occupancy on the structure of the generated clusters and their similarity to real protein structure.

To investigate the effect of occupancy, we use the random relaxation procedure because there is very little difference between the random move algorithm and the more computationally demanding RMC–SA algorithm. Three occupancies are studied: 20, 40 and 60%. In the case of BOO parameters, increasing the occupancy led to the narrowing of the *Q*_*l,i*_ distributions and shifted them towards lower mean values (Figure 9 and Table 3). For *l* = 2 and 6, *Q*_*l,i*_ distributions of the generated structures at 40% occupancy matched the protein distribution more closely than those at 20 and 60% occupancies. However, for *l* = 4, 60% occupancy distribution is the closest to that of the protein. These observations are also reflected in the Kolmogorov–Smirnov statistics (Table 4). The BOO parameter *Ŵ*_*l,i*_ is not sensitive to occupancy especially for *l* = 4 and 6. For example, the width of the *Ŵ*_*l,i*_ distributions do not change, in fact, the width do not change for any *l*. On the other hand, for *l* = 2, the effect of occupancy is quite complex. The average protein structure shows an almost flat distribution around *Ŵ*_2,*i*_ = 0 with a slight positive slope. On the other hand, the generated structures with 20% occupancy show a large negative slope. Increasing the occupancy increased the slope around *Ŵ*_2,*i*_ = 0. At 60% occupancy, the *Ŵ*_2,*i*_ distribution of the generated structures become almost flat and is the closest to that of the average protein. This observation is also supported by the Kolmogorov–Smirnov analysis, which also indicates that the 40% and 60% occupancies are statistically identical to each other and to the average protein (Table 4).

**Table 3.** The influence of occupancy on the mean values of various Bond Orientational Order and network parameters for unrelaxed on–lattice clusters and clusters relaxed with random move algorithm. The average eigenvalue of the normalized Laplacian is not shown because it is always equal to one by definition. Standard error is shown within parenthesis for the last digit and only when it is greater than 1%.

| Parameter | Protein | Occupancy |  |  |  |  |  |
| --- | --- | --- | --- | --- | --- | --- | --- |
|  |  | 20% |  | 40% |  | 60% |  |
|  |  | Unrelaxed<br>(On–<br>lattice) | Relaxed<br>(Random) | Unrelaxed<br>(On–<br>lattice) | Relaxed<br>(Random) | Unrelaxed<br>(On–<br>lattice) | Relaxed<br>(Random) |
| $\bar{Q}_{2,i}$ | 0.331 | 0.496 | 0.498 | 0.343 | 0.352 | 0.233 | 0.245 |
| $\bar{Q}_{4,i}$ | 0.270 | 0.419 | 0.441 | 0.310 | 0.334 | 0.213 | 0.237 |
| $\bar{Q}_{6,i}$ | 0.345 | 0.434 | 0.455 | 0.317 | 0.349 | 0.215 | 0.260 |
| $\overline{\hat{W}}_{2,i}$ | 0.012 | -0.071 | -0.071 | -0.028 | -0.030 | 0.005 | -0.004 |
| $\overline{\hat{W}}_{4,i}$ | 0.016 | 0.005 | 0.024 | 0.005 | 0.015 | -0.001 | 0.008 |
| $\overline{\hat{W}}_{6,i}$ | -0.004 | -0.018 | -0.025 | -0.009 | -0.017 | 0.003 | -0.009 |
| $\bar{k}$ | 7.947 | 4.811 | 4.717 | 7.381 | 7.162 | 11.275 | 10.919 |
| $\bar{k}_{nn}$ | 8.344 | 5.330 | 5.204 | 8.080 | 7.799 | 12.181 | 11.732 |
| $\bar{C}$ | 0.550 | 0.618 | 0.605 | 0.547 | 0.542 | 0.530 | 0.528 |
| $\bar{L}$ | 5.129 | 5.348 | 5.394 | 4.199 | 4.225 | 3.921 | 3.850 |

**Table 4.** The influence of the occupancy on the two–sample Kolmogorov–Smirnov test parameters for structures relaxed with the random move algorithm. The on–lattice and RMC–SA relaxed results follow the same trends observed in the randomly relaxed structures. Italic text indicates *p*– values less than 0.05.

| Parameter | Occupancy |  |  |  |  |  |
| --- | --- | --- | --- | --- | --- | --- |
|  | 20% |  | 40% |  | 60% |  |
| | $D$ | $p$-value | $D$ | $p$-value | $D$ | $p$-value |
| $Q_{2,i}$ | 0.488 | *0.011* | 0.123 | 0.996 | 0.389 | 0.071 |
| $Q_{4,i}$ | 0.481 | 0.012 | 0.242 | 0.539 | 0.152 | 0.961 |
| $Q_{6,i}$ | 0.381 | 0.083 | 0.116 | 0.998 | 0.460 | 0.019 |
| $\hat{W}_{2,i}$ | 0.243 | 0.376 | 0.123 | 0.983 | 0.049 | 1.000 |
| $\hat{W}_{4,i}$ | 0.052 | 1.000 | 0.011 | 1.000 | 0.046 | 1.000 |
| $\hat{W}_{6,i}$ | 0.221 | 0.493 | 0.134 | 0.961 | 0.073 | 1.000 |
| $k_i$ | 0.544 | 0.002 | 0.154 | 0.937 | 0.370 | 0.074 |
| $k_{nn,i}$ | 0.747 | 0.000 | 0.176 | 0.848 | 0.673 | 0.000 |
| $C_i$ | 0.118 | 1.000 | 0.127 | 1.000 | 0.088 | 1.000 |
| $L_i$ | 0.114 | 0.997 | 0.276 | 0.319 | 0.440 | 0.019 |
| $\lambda$ | 0.148 | 0.975 | 0.049 | 1.000 | 0.082 | 1.000 |

**Figure 9.**
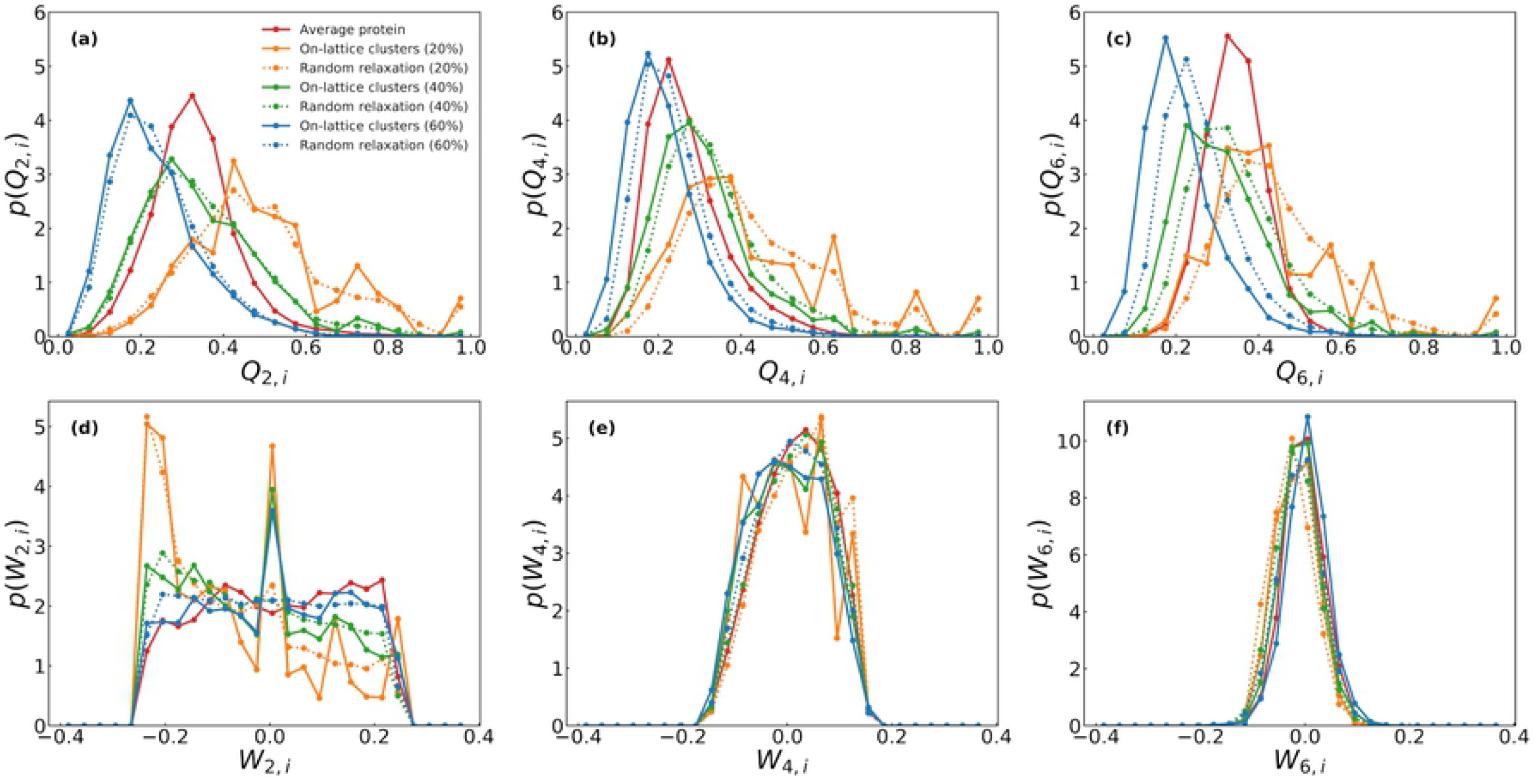
The influence of occupancy on various BOO parameters for *l* =2, 4, and 6.

The degree and average neighbor degree show similar trends with respect to occupancy; the distributions of the generated structures become wider and shift towards greater mean values Figure 10(a) and (b)). In both cases, 40% occupancy is the closest to the average protein. In fact, at 20 and 60% occupancy, the degree and nearest neighbor degree distributions of the generated structures show no statistical similarity to that of the average protein as suggested by the Kolmogorov–Smirnov analysis (Table 4). The relationship between the degree and occupancy is quite straightforward; the number of neighbors for any lattice site increases as a result of increasing occupancy. The same effect is also reflected in the nearest neighbor degree. It is obvious that a larger occupancy than 40%, but less than 60% is required to match the degree and average neighbor degree distributions of real proteins.

**Figure 10.**
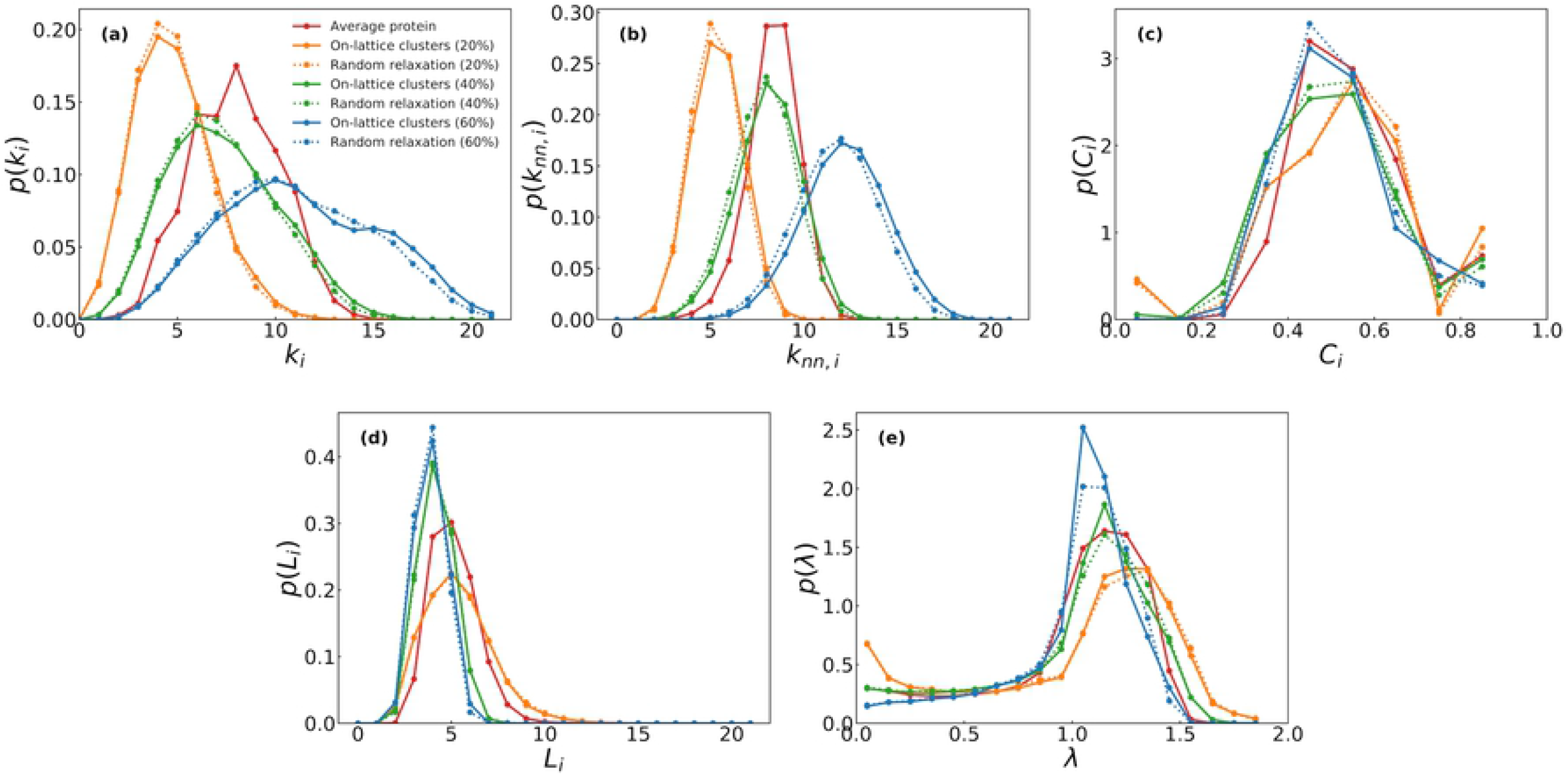
Comparison of the distributions of (a) degree (contact number), (b) nearest neighbor degree, (c) clustering coefficient, (d) the shortest path length, and (e) the eigenvalue of the normalized Laplacian for the average protein and generated clusters with respect to cluster occupancy.

The clustering coefficient distributions show slight variations with respect to different percentage of occupancy; the distributions become slightly narrower with lower mean values upon increasing occupancy (Figure 10(c)). The clustering coefficient distribution at 60% occupancy is closest to that of the average protein, although the Kolmogorov–Smirnov analysis did not reveal any statistical difference with respect to occupancy. This observation shows that a protein residue on average has a more densely connected neighborhood than the cubic cluster, a characteristic which is not fully governed by a global occupancy parameter that is used in the current study.

In the case of the shortest path length, the distributions of the generated structures shift towards lower mean values and become narrower with increasing occupancy (Figure 10(d)). This is expected given that at greater occupancy, there is a greater possibility of finding a shorter path between any two residues. Examination of the Laplacian eigenvalue distributions show that the number of vertex doubling and the number of connected components increased with increasing occupancy. This is probably due to the connectivity constraint used during emptying of the lattice clusters which leads to a branched structure.

## 4 Conclusions

In this work, we contribute to the toolbox of methodologies aiming at characterizing as well as generating artificial structures that automatically reproduce the general structural characteristics of globular proteins. We show that simple cubic (SC) lattice is the proper structural template for such a construction. The relevance of cubic lattices has been originally motivated from the statistical distributions of internal coordinates defined by C_α_ atoms (i.e., C_α_–C_α_–C_α_ bending angle) (36). In the present study, we show that defect laden SC arrangement is the proper cubic template and the closed packed FCC is incompatible with residue coordination characteristics. This observation is directly motivated from the average protein radial distribution function (RDF).

We identify three major sources of disorder that are sufficient to recover the geometric and topological properties of proteins with an artificial network structure if we start from a bulk SC crystal: The molecular (free) surface, random distribution of voids, and small displacements from ideal lattice positions.

Molecular surface accounts for the finite size of protein molecules and is an obvious requirement. Cubic clusters are easy to generate and can conveniently replicate the compact shape characteristics of globular proteins. This choice is justified *a posteriori* as the shortest path lengths of generated structures match well with that of real proteins.

Our results single out randomly distributed voids as the primary source of disorder needed to reproduce the structural metrics of globular proteins. Most notably, the mean coordination number in the SC cluster mimics protein characteristics at an arbitrary cut-off only after introducing voids at a concentration of 60% into a perfect cluster. This observation is the core of our structure generation strategy. Since the voids are introduced in accordance with general packing trends with no specific structure in mind they do not have direct structural or functional interpretation. However, the evident ability to generate a large number of representative structures at once simply by shuffling point defects at a predetermined concentration is noteworthy. Considering that BOO parameters, which are energy-like quantities, are largely unaffected by this redistribution of voids hints that the globular protein landscape might be inherently degenerate at this level of coarse-graining.

A small positional disorder is needed to recover the necessary spread around the peaks that are present in the protein RDF, but these displacements do not disrupt other properties. Unlike simple liquids, the amount of positional disorder that needs to be introduced is rather limited. The combined reverse Monte Carlo and simulated annealing (RMC–SA) procedure recovers the average RDF of 210 proteins by perturbing the on-lattice residues of a finite sized structure with voids. The magnitude of the displacements produced by the reconstruction procedure do not significantly differ for the surface or core residues and stays well below the nearest neighbor distance (∼3.8 Å) of proteins. Neither bond orientational order parameters (BOO) nor the network characteristics are affected by the RMC–SA simulations. All the resulting structures are shown to nontrivially approximate several topological and geometrical features of real proteins. Therefore, the correspondence between the defect laden SC clusters and real residue networks is not superficial and positional disorder is not a critical ingredient. Our approach is novel and uniquely general in the sense that it enables ensemble-versus-ensemble comparisons without targeting individual protein structures.

We, however, note that the structure generation method does not ensure that the resulting structures form self-avoiding walks through nearest neighbor contacts. One possible strategy for ensuring self-avoiding walk property is to deepen the crystal lattice analogy to introduce extended defects. Most notably, a polycrystalline grain structure can be deployed wherein our ground rules could be imposed to generate smaller fragments. Such smaller self-avoiding fragments generated on each grain would then be tied together by reorienting each grain to yield the final self-avoiding polymer. However, to realize this, we need a set of well-defined rules to determine the optimal size of the grains and a meaningful texture map (orientation distributions of the grains), which we currently lack.

Finally, there has been a sustained interest in the computational manipulation of targeted self–assembly (75,76) and building simple pair potentials that produce various open structures with controlled defect concentration as their ground state (77,78). Such a directed self–assembly approach is not readily applicable to proteins as it is hard to define candidate ground states with simple rules. However, culling novel tools from inverse optimization efforts might be a viable alternative to the widely used statistical potentials. The current work, by singling out SC arrangement as a candidate background and presenting a physically motivated recipe for structure generation, is a step forward in this direction.

## 5 Acknowledgements

This material is partially based upon the work supported by the National Science Foundation (NSF) under Grant Nos. 1538730 and 1825254 and the Scientific and Technological Research Council of Turkey (TUBITAK) under Grant No. 117F389. The authors acknowledge computational support provided by the Rensselaer Center for Computational Innovations.

## Footnotes

† For spherical harmonics, we adopted the standard definition and ignored the Condon-Shortley phase (–1)^m^, which leads to 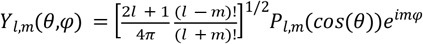, where *P*_*l,m*_ (*cos*(*θ*)) are the associated Legendre functions. Normalization constant is consistent with Steinhardt et al. (50) and yields 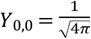.

